# BEADS: An Interactive Semi-Automated Workflow for 3D Fibrin Angiogenesis Assays Enabling Co-Culture and Directionality Analysis

**DOI:** 10.64898/2025.12.16.694634

**Authors:** Cailin R. Gonyea, Brielle Hayward-Piatkovskyi, Jason P. Gleghorn

## Abstract

3D fibrin bead angiogenesis assays are widely used to study endothelial sprouting *in vitro*, yet current analytical approaches are either time-consuming or poorly adaptable to complex imaging conditions, limiting quantitative assessment of co-cultures, spatial interactions, and nearest-neighbor–dependent angiogenic behavior. In this study, we developed a semi-automated user-interactive image analysis pipeline, Bead-based Endothelial Angiogenesis Data Suite (BEADS), to provide standardized quantitative bead-centric metrics of sprouting, migration, and spatial orientation in 3D fibrin angiogenesis assays. BEADS integrates automated bead detection with manual correction, followed by guided sprout and migratory-cell annotation across multi-channel image z-stacks. Novel analytical capabilities include co-culture designation, nearest-neighbor pairing, and circular statistics for sprout-directionality quantification. Performance was evaluated in assays using co-cultured male and female human pulmonary microvascular endothelial cell (HPMEC)-coated beads. BEADS reduced hands-on analysis time approximately sevenfold compared with manual tracing while preserving sprout-length accuracy against manual ground truth. BEADS provides a standardized, extensible platform for microvascular image analysis, supporting co-culture experimentation, spatial endothelial-interaction metrics, migratory-cell quantification, and high-throughput adaptation. This semi-automated workflow enables quantitative microvascular research by integrating computational precision with endothelial behavior and is broadly applicable to angiogenesis assays that incorporate co-cultures, perturbations, or multi-label experimental designs.

## INTRODUCTION

Angiogenesis, the formation of new blood vessels from pre-existing vasculature, is a fundamental process in both physiological tissue maintenance and pathological remodeling (1–7). Understanding the microvascular mechanisms that regulate endothelial sprouting, migration, and spatial coordination remains critical for developing therapies across cancer, ischemia, inflammation, and tissue engineering (1, 8–13). Quantitative *in vitro* models play an essential role in uncovering these mechanisms, particularly those capable of capturing behaviors at the single-sprout level, where microvascular patterning emerges.

The 3D fibrin bead angiogenesis assay is a well-established *in vitro* model that recapitulates multicellular sprouting, endothelial migration, and matrix invasion under controlled biochemical and mechanical cues (**Fig. 1**) (14–17). In this assay, endothelial cells are coated on the surface of microcarrier beads that are subsequently embedded in fibrin gels, producing radially oriented sprouts that mimic capillary-scale morphogenesis. The assay provides a controlled environment for investigating the factors that influence angiogenesis, including cell–cell interactions and soluble and physical extracellular matrix (ECM) cues. Metrics such as sprout number, sprout length, and migratory-cell behavior are routinely used to assess angiogenic potential.

**Fig. 1.**
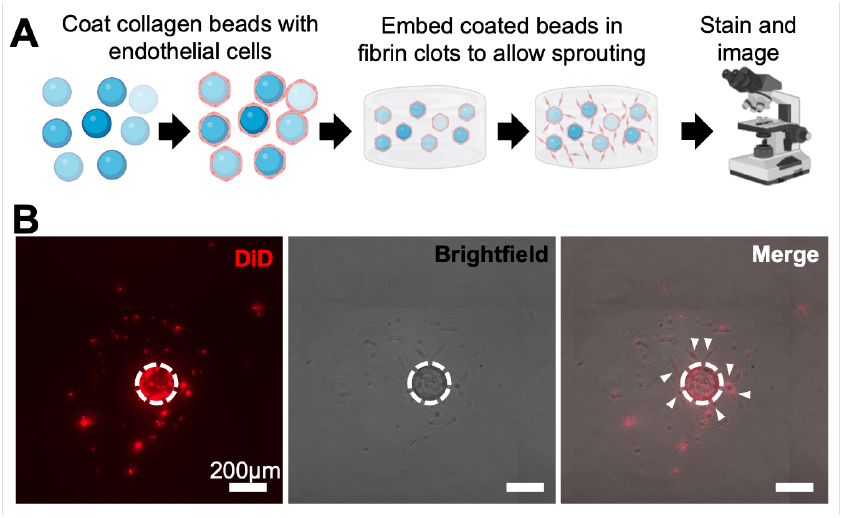
Fibrin 3D bead angiogenesis assay generates complex imaging outputs that challenge fully automated analysis. **(A)** Schematic of the assay workflow, in which collagen microcarrier beads coated with endothelial cells, are embedded within fibrin gels, and undergo multicellular sprouting prior to imaging. **(B)** Representative fluorescence and brightfield images showing endothelial sprouts (arrows) extending from beads (dotted outline). These heterogeneous imaging conditions—including variable sprout density and mosaic fluorescence underscore the need for semi-automated, user-guided analysis pipelines such as BEADS. Scale bar = 200 *µ*m.

However, existing analysis tools, including ImageJ-based pipelines, intensity-thresholded segmentation approaches, and recent machine-learning frameworks (18), remain limited for bead-centric assays. These pipelines generally perform poorly on heterogeneous fluorescence, do not support co-culture discrimination, and cannot compute nearest-neighbor–dependent metrics, which are increasingly recognized as important drivers of microvascular pattern alignment (19–23). A further limitation of current tools is that fully automated segmentation often fails under realistic experimental conditions including overlapping sprouts, variable dye intensity, mosaic staining, or dense co-cultures, leading to either misidentification or dataset exclusion. Conversely, manual tracing is accurate but prohibitively slow for multi-well studies or screening applications. The field, therefore, lacks an approach that balances automation, accuracy, and user oversight, particularly for assays interrogating cell–cell interactions such as sex differences, donor variability, or mixed endothelial phenotypes.

To address these unmet needs, we developed the Bead-based Endothelial Angiogenesis Data Suite (BEADS), a semi-automated, user-interactive workflow for quantitative analysis of 3D fibrin angiogenesis assays. BEADS is specifically designed to (1) identify beads with automated algorithms and user-guided correction, (2) quantify sprout number and length, (3) track migratory cells, (4) label beads by co-culture identity, and (5) compute nearest-neighbor relationships and directionality metrics using circular statistics. Together, these capabilities enable quantitative questions that were previously inaccessible using existing tools. Here, we apply BEADS to a representative co-culture model using male and female human pulmonary microvascular endothelial cells (HPMECs) to validate the pipeline and demonstrate its ability to resolve sex-dependent migratory behavior and nearest-neighbor–dependent sprout orientation. This framework provides a standardized, extensible platform for microvascular image analysis and supports the broader goal of quantifying endothelial interactions and coordination underlying microvascular patterning.

## MATERIALS AND METHODS

### Cell Culture

Human pulmonary microvascular endothelial cells (HPMECs, ScienCell, Carlsbad, CA; female lots: 17799, 17806, 15900; male lots: 11816, 11422, 16021) were cultured as previously described (17) on fibronectin-coated plates (2 *µ*g/cm^2^) in hormone-free media (HFM) and grown in phenol red–free endothelial cell medium (ScienCell) at 37°C supplemented with 5% CO_2_. HFM was supplemented with endothelial cell growth supplement (ScienCell), 1% penicillin-streptomycin (ScienCell), and 5% charcoal-stripped FBS (HyClone, Logan, UT). Individual donors were pooled by sex to reduce donor-to-donor variability, generating male- or female-pooled HPMECs, which were then grown to near confluence before experimental use. Cells were labeled in HFM media with 1 *µ*M/10 mL lipophilic carbocyanine dyes (Biotium, Fremont, CA), DiO (M) or DiD (F), to differentiate the sex of the cells. Cells were incubated in dye for 20 min at 37°C per manufacturer’s instructions. HPMECs were used between passages 4 to 6. All endothelial cell lots were confirmed for CD31^+^/VE-cadherin^+^ phenotype to ensure microvascular identity.

### 3D Fibrin Bead Angiogenesis Assay

Collagen-coated Cytodex-3 microcarrier beads (Sigma-Aldrich, St. Louis, MO) were (**?**) coated with male or female HPMECs at 20,000 cells per 750 beads, incubated for 4 h at 37°C with periodic agitation, then statically overnight, as previously described (16, 24). Male and female beads were pooled and resuspended in 2 mg/mL fibrin (Millipore, Burlington, MA) gels supplemented with 0.15 U/mL of aprotinin (Sigma-Aldrich) at 250 beads/mL. Gel cultures were maintained in HFM for 3 days with daily medium changes. After 3 days, wells were imaged using a Zeiss epifluorescent microscope at 10x magnification (**Fig. 1A**). Sprouting within this period produced multicellular extensions of 50-150 *µ*m, consistent with capillary microvessel dimensions and previous microvascular morphometry (**Fig. 1B**). To image each well, 81 (9×9) image tiles with 48 z-slices spaced 3 *µ*m apart were used.

### Image Analysis Using Bead-based Endothelial Angiogenesis Data Suite (BEADS)

MATLAB version 9.11.0.1769968 (R2021b) was used for the development of BEADS. Image segmentation parameters (e.g., bead radius range, threshold values) were empirically optimized to maintain inter-well consistency and minimize variation in bead detection. Images were analyzed for the following characteristics: percentage of beads per sex, percentage of beads with at least one sprout, average number of sprouts per bead, maximum sprout length, average sprout length, and normalized number of migratory cells per well.

### Image Analysis Comparison Between BEADS, Manual Hand-Tracing and the ImageJ Angiogenesis Analyzer

To compare our semi-automated method to established standards, images of the same 88 beads (regardless of sex) across three wells were analyzed using BEADS, manual hand tracing, and the ImageJ Angiogenesis Analyzer. Sprouts were identified, and the length of each sprout, measured from the tip to the nearest point on the bead edge, was recorded. The raw sprout-length data were then processed either manually or using the post-processing BEADS MATLAB script to calculate the maximum and mean sprout lengths per bead for comparison. The analysis time for each well was recorded and normalized to the number of beads. Timing included sprout selection and data export; ground truth for length was the straight-line centroid-to-tip distance (bead radius subtracted) for both methods to ensure comparability.

### Radial Histograms/Circular Statistics

After analysis using BEADS, angiogenic beads were sorted into the following categories based on their sex and the sex of the nearest neighbor: male–male, male–female, female–male, female–female. Angles between each bead’s centroid and its nearest neighbor were calculated using the coordinates of both the bead center and sprout tip. The angles were then normalized so that the nearest neighbor was oriented at 0°. This was repeated for every bead in each category. Radial histograms were constructed using MATLAB *polarhistogram*, and circular statistics were computed using the MATLAB *CircStat* package (25). Because the data appeared multimodal, we performed the Rao spacing test to determine uniformity (26), followed by a V-test with a specified mean direction of 0° to evaluate whether sprouting was directionally biased toward the nearest neighbor (NN). This analysis was used to determine the effects of NN sex on sprouting directionality. All circular statistical analyses were conducted using MATLAB version 9.11.0.1769968 (R2021b).

### Statistical Analysis

Data from BEADS were used to report the percentage of each sex within a well, the percentage of beads with at least one sprout per well, the average number of sprouts per bead, the average sprout length, the maximum sprout length, and the number of migratory cells for both males and females. All groups were assessed for normality using a Shapiro–Wilk test for groups with fewer than eight biological samples or using a D’Agostino–Pearson omnibus test for groups with eight or more samples. If normally distributed, an unpaired Student’s t-test was used; otherwise, a Mann–Whitney test was applied. Values of *p <* 0.05 were considered statistically significant. All statistical comparisons were performed using GraphPad Prism version 9.5.1 (GraphPad Software, Boston, MA). All data are represented as mean *±* standard deviation (SD) or mean *±* interquartile range for violin plots.

## RESULTS

### A Semi-Automated Computational Framework for Quantifying 3D Microvascular Sprouting

We first confirmed that the fibrin bead assay produced microvessel-like multicellular sprouts with continuous endothelial morphology and expected branching behavior (**Fig. 1B**). BEADS was then applied to these datasets to generate bead-centric and sprout-level quantitative outputs. The full semi-automated workflow is described below.

#### Adaptive Image Processing for Reliable Well Segmentation and Bead Detection

The BEADS pipeline begins with multichannel z-stacks that include brightfield (BF) and fluorescence channels. The central slice from the brightfield channel z-stack of images is extracted **(Fig 2Ai.)** and a circular Hough transform is applied to identify the well boundary in the well plate and pixels outside this boundary are masked **(Fig 2Aii.)**. Large circular artifacts (e.g. bubbles) are removed via a second Hough detection using radii above the expected bead range to remove them from analysis **(Fig 2Aiii.)**. A Laplacian-of-Gaussian filter smooths the image **(Fig 2Aiv.)**, followed by image binarization and subsequent Hough-based bead detection **(Fig 2Av.)**. Bead centroid coordinates (x,y) and radii are recorded. After identification of the beads, the brightfield image is displayed in a graphical user interface (GUI) with editable circular ROI handles overlaid onto each identified bead **(Fig 2Avi.)**. Users can add missed beads **(Fig 2Bi.)**, adjust bead size/position, or remove misidentified ROIs via the GUI **(Fig 2Bii.)**. Automated detection captured ~92–96% of beads, and the remaining beads were user-corrected. BEADS pauses until the user closes the GUI, ensuring complete bead-set verification.

**Fig. 2.**
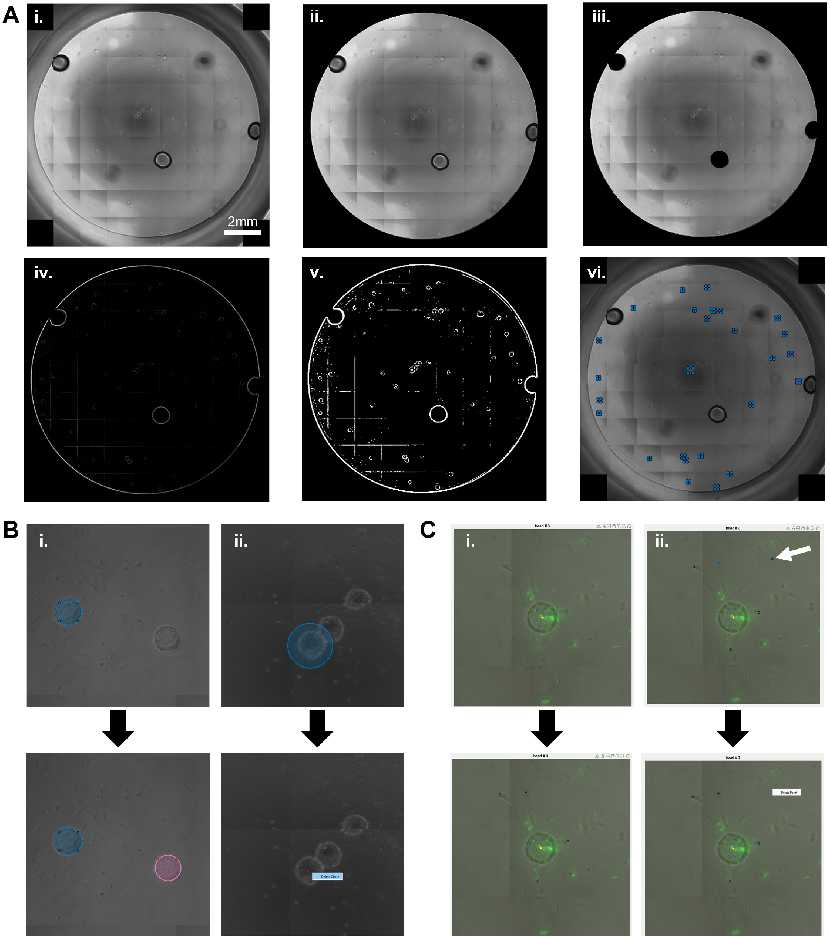
Semi-automated BEADS workflow enables reliable bead detection and user-guided correction prior to sprout annotation. **(A)** Automated preprocessing includes extraction of the central brightfield slice (i), masking of the well boundary (ii), removal of large circular artifacts such as bubbles (iii), image smoothing with a Laplacian-of-Gaussian filter (iv) and binarization with Hough-based bead detection (v). Detected beads are presented in a GUI with editable ROI handles (blue points) (vi). **B)** Users can add missed beads (i, pink ROI) or delete misidentified objects (ii) to ensure accurate bead representation. **C)** For each bead, sprout tips (blue dots) are selected via mouse click (i), with right-click removal of erroneous points (ii). Users then assign a population label (“A” or “B”) before BEADS advances to the next bead. This combination of automated detection and guided correction ensures accurate identification of beads and sprouts across heterogeneous imaging conditions.

#### Interactive Sprout Mapping Enables High-Resolution Microvascular Phenotyping

Once the bead ROIs are finalized, the brightfield image is concatenated with the maximum-intensity projections of any fluorescent images, and the sprout-annotation GUI presents the merged BF and fluorescent image to visualize the beads with the sprouts **(Fig 2C)**. Users click to mark (x,y) coordinate of sprout tips **(Fig 2Ci.)**, and right-click to remove mislabeled points **(Fig 2Cii.)**. After identifying all sprouts, users can classify the bead as belonging to a population A or B if multiple cell populations or phenotypes are being tracked, enabling the detection of within- and between-population (group) differences. This annotation structure supports bead-level metadata retention, which is critical for nearest-neighbor and population-dependent analyses.

#### Semi-Automated Detection of Detached Migratory Endothelial Cells

Migratory cells, defined as cells detached from beads or sprouts, were annotated similarly to sprout tip selection, except that the user selects migratory cells within the GUI instead of spout tips. The individual bead images are presented sequentially for annotation of migratory cells. If there is multiple population tracking, users click all migratory cells belonging to population A, then repeat the bead cycle for population B. Counts are normalized per bead to allow comparison across wells.

#### Standardized Extraction of Sprout Metrics and Spatial Interaction Parameters

BEADS generates separate bead- and sprout-data tables containing centroids, radii, sprout coordinates, and population identity. These files are exported as text files and are concatenated into a single Excel sheet for data processing. Similarly, migratory cells are stored in a table with columns for the population A count and the population B count. This data table is then also exported into an Excel file. The BEADS post-processing scripts calculate the number of sprouts per bead and the mean and maximum sprout lengths by measuring each sprout’s length as the straight-line distance from the tip (x, y) coordinates to the bead centroid, then subtracting the bead radius. BEADS also assesses each bead’s centroid coordinates and identifies the bead with the closest centroid coordinates to determine the nearest neighbor (NN) bead. After identifying the NN, the script reports the index of each bead’s NN, the number of sprouts, and the population (A or B) of the NN, as well as the distance to the NN in pixels. These outputs enable analysis of sprout magnitude, distribution, and spatial orientation, providing insights inaccessible to traditional tools.

### Sex-Dependent Angiogenic and Migratory Behavior in Co-Cultured HPMECs

To validate BEADS, we applied it to quantify angiogenic behavior in co-cultured male (M) and female (F) HPMEC-coated beads within the same fibrin environment **(Fig 3A-B)**. This design enabled a direct comparison of endothelial performance under identical microenvironmental conditions. Sex-dependent endothelial behavior, including differences in motility, proliferative capacity, and paracrine signaling, has gained increasing attention in microvascular biology (27–29), making co-culture assays well suited for identifying interaction-dependent phenotypes.

**Fig. 3.**
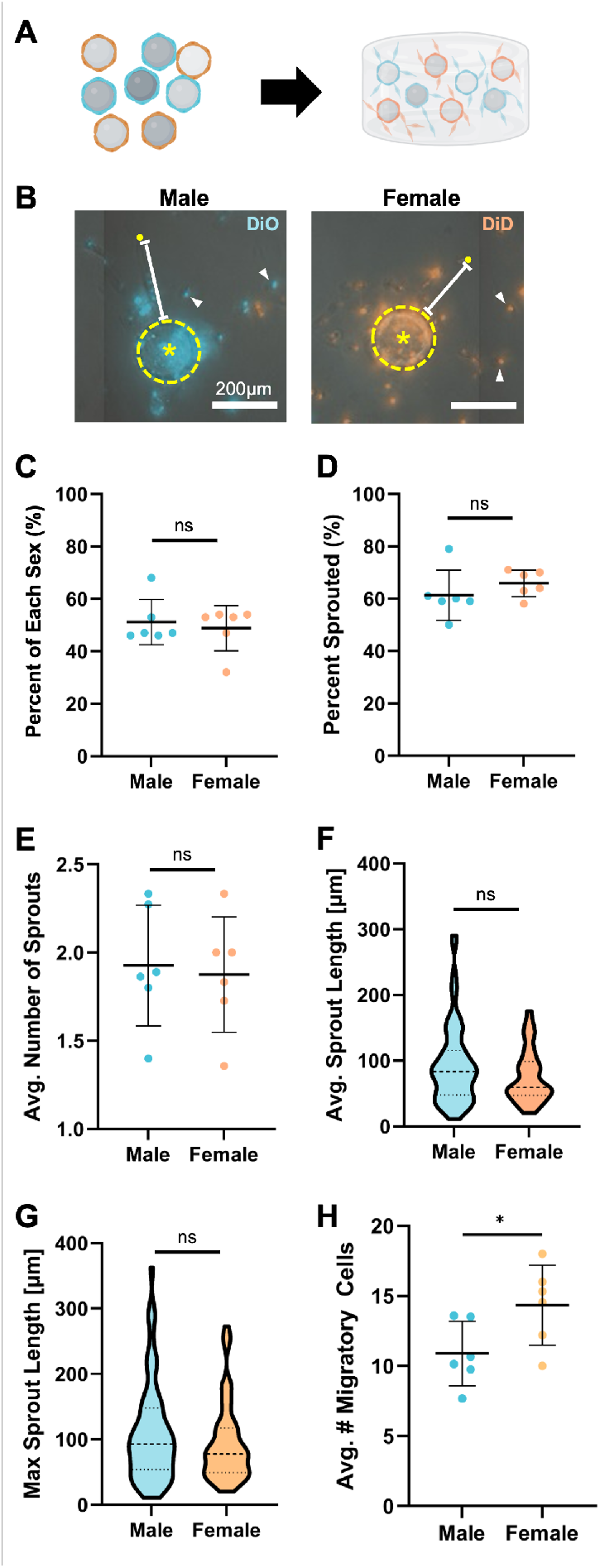
BEADS quantifies sprouting and migratory behavior in co-cultured male (M) and female (F) HPMEC-coated beads. **A)** Experimental design in which M (blue) and F (orange) HPMEC-coated beads are pooled and embedded together in fibrin and co-cultured for 3 days. **B)** Representative images showing beads (yellow dashed outline), bead centroid (yellow asterisk), sprout tips (yellow dots), and migratory cells (white arrows). Scale bar = 200 *µ*m. Quantification of angiogenic metrics, including the proportion of beads of each sex per well **C)**, percentage of beads producing ≥1 sprout **D)**, number of sprouts per bead **E)**, mean sprout length **F)**, and maximum sprout length **G)**. No significant sex-dependent differences were observed in these sprouting parameters. **H)** Female-coated beads exhibited significantly more migratory cells per bead than male-coated beads (p = 0.0433), consistent with sex-dependent endothelial motility. Data represent n = 6 wells for C-E; n = 68 M and 65 F beads for F-G; violin plots show mean ± IQR; dot plots show mean ± SD.

Across six wells, BEADS detected an equivalent distribution of M and F beads (51.57% vs. 48.83%, p = 0.5887), confirming uniform mixing upon gel preparation **(Fig 3C)**. The percentage of beads that produced at least one sprout was similar between sexes (61.33% M vs. 65.81% F, p = 0.3303) **(Fig 3D)**, and the average number of sprouts per bead did not differ between sexes (1.926 ± 0.3417µm vs. 1.875 ± 0.3265µm, p = 0.7959) **(Fig 3E)**. Mean sprout length (92.46 ± 59.01µm vs. 75.34 ± 39.42µm, p = 0.9576) **(Fig 3F)** and maximum sprout length (108.7 ± 74.55µm vs. 93.43 ± 59.09µm, p = 0.2964) **(Fig 3G)** were likewise comparable between M and F beads. Together, these findings indicate that male and female HPMECs exhibit similar intrinsic sprouting capacity in hormone-free medium, validating the use of this model for probing interaction-driven differences in endothelial behavior.

In contrast, BEADS revealed a significant difference in endothelial migration. Migratory cells—defined as cells detached from bead or sprout structures were identified and normalized to bead number **(Fig 3B, arrowhead**). Female-coated beads produced more migratory cells per bead than male-coated beads (14.35 ± 2.849 vs. 10.89 ± 2.297; p = 0.0433) **(Fig 3H)**. These results indicate the enhanced migration of female HPMECs and mirrors sex-dependent endothelial motility observed in microvascular wound healing models(16, 17, 30–32). Together, these validation studies demonstrate BEADS’s sensitivity to detecting subtle endothelial behavioral phenotypes that may be obscured using whole-well metrics alone(17, 32).

### Directional Sprouting is Influenced by Nearest-Neighbor Sex

To determine whether nearby endothelial populations influence the spatial orientation of sprouting, BEADS assigned each bead a nearest neighbor (NN) based on centroid proximity and computed sprout angles relative to the bead-NN axis **(Fig 4AB)**. Sprout-angle distributions were significantly non-uniform across all NN groupings (p < 0.001), demonstrating anisotropic sprouting behavior. No significant directional bias toward the NN (0°) was found for M–M, F–F, or F–M pairs (p = 0.9484, 0.3490, and 0.0987, respectively) **(Fig 4C)**. However, M–F pairs showed a significant orientation of sprouting toward the female NN (p = 0.0007). No group exhibited preferential sprouting away from the NN (180°). These findings demonstrate that female endothelial neighbors induce directional sprouting from male-coated beads, suggesting short-range paracrine or juxtacrine influences on microvascular pattern alignment. Such relational metrics have not been accessible through existing angiogenesis analysis approaches, underscoring the unique analytical capabilities of BEADS.

**Fig. 4.**
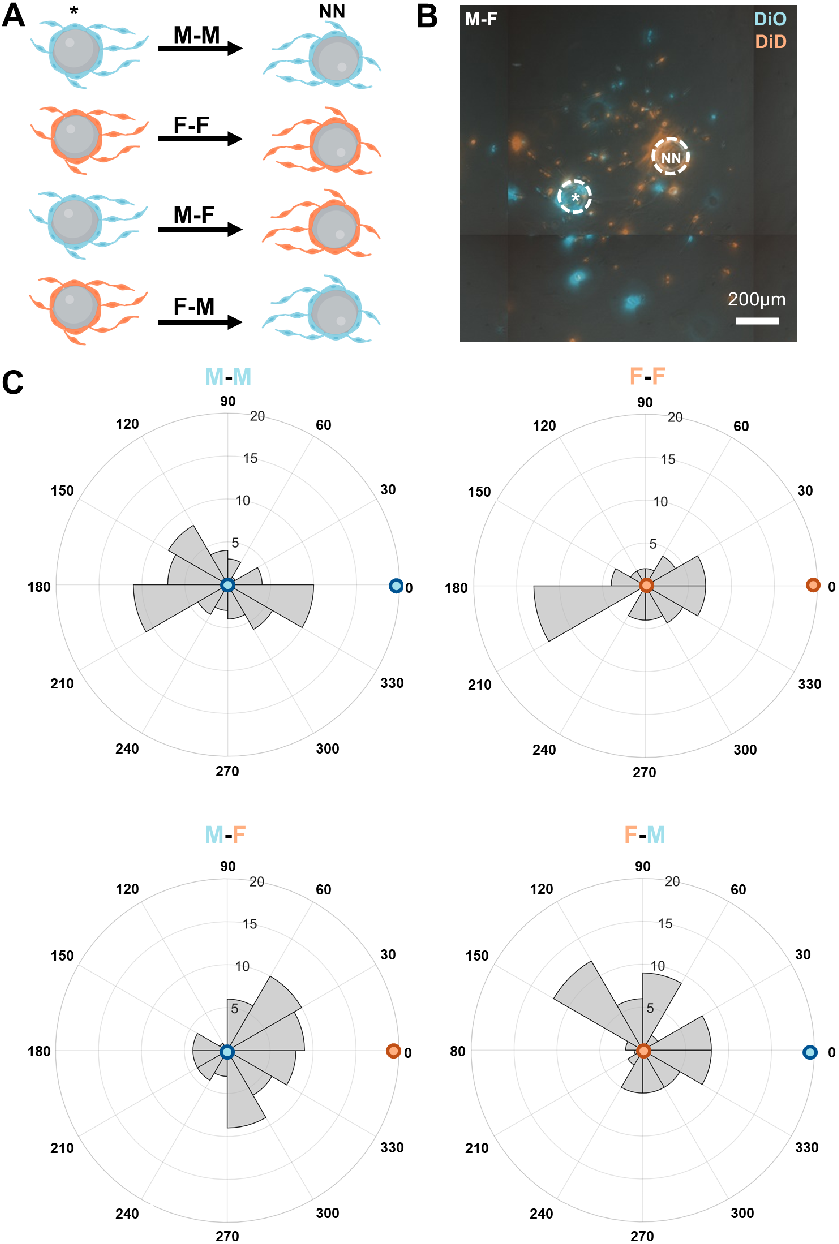
BEADS reveals nearest-neighbor–dependent directional sprouting in co-cultured endothelial populations. **A)** Schematic showing the four nearest-neighbor (NN) pairings evaluated (M–M, F–F, M–F, and F–M) where the first bead is the bead of interest (asterisk) and the second bead is the NN. **B)** Representative fluorescence image of co-cultured beads (white dashed outlines) used for directional analysis. Scale bar = 200 *µ*m. **C)** Radial histograms showing sprout-angle distributions normalized such that the NN lies at 0°. Male beads with female NNs (M–F) exhibited significant directional bias toward the NN (V-test p = 0.0007), whereas other pairings showed no significant directional preference. Blue and orange dots indicate male and female populations, respectively. M-M n = 48 beads, 65 sprouts; F-F n = 42 beads, 54 sprouts; M-F n = 59 beads, 64 sprouts; F-M n = 57 beads, 65 sprouts.

### Comparison with Manual and Automated Analysis Tools

To validate the performance of the BEADS, its sprout-length measurements and annotation times were compared against manual hand-tracing and the ImageJ Angiogenesis Analyzer plugin. The ImageJ plugin was included for comparison but it could not be quantitatively compared because it consistently failed to identify beads and sprouts under co-culture and mosaic-staining conditions **(Fig 5)**. Across 88 beads from three wells, BEADS reduced hands-on annotation time by approximately sevenfold relative to manual tracing (p = 0.0010) while maintaining accuracy for both average and maximum sprout length measurements **(Fig 6)**. This substantial increase in throughput, combined with preserved accuracy and user-interactive correction, highlights the practicality of BEADS for large-scale angiogenesis studies, including longitudinal or pharmacologic assays.

**Fig. 5.**
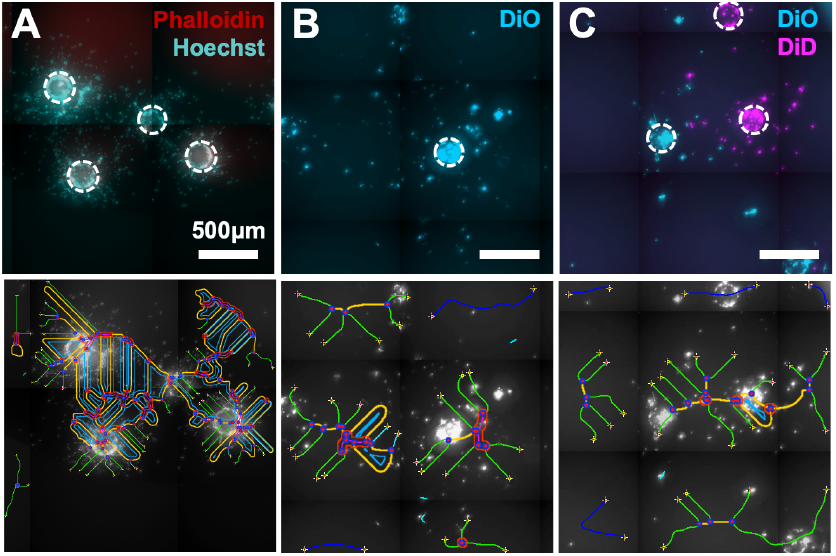
Fully automated analysis fails under common imaging challenges encountered in 3D fibrin angiogenesis assays. Representative examples of conditions that disrupt automated segmentation, including high cell density **(A)**, mosaic fluorescence staining **(B)**, and multi-population co-cultures **(C)**. Lower panels show the result of fully automated algorithms frequently misidentify beads and sprouts under these conditions. Scale bars = 500 *µ*m.

**Fig. 6.**
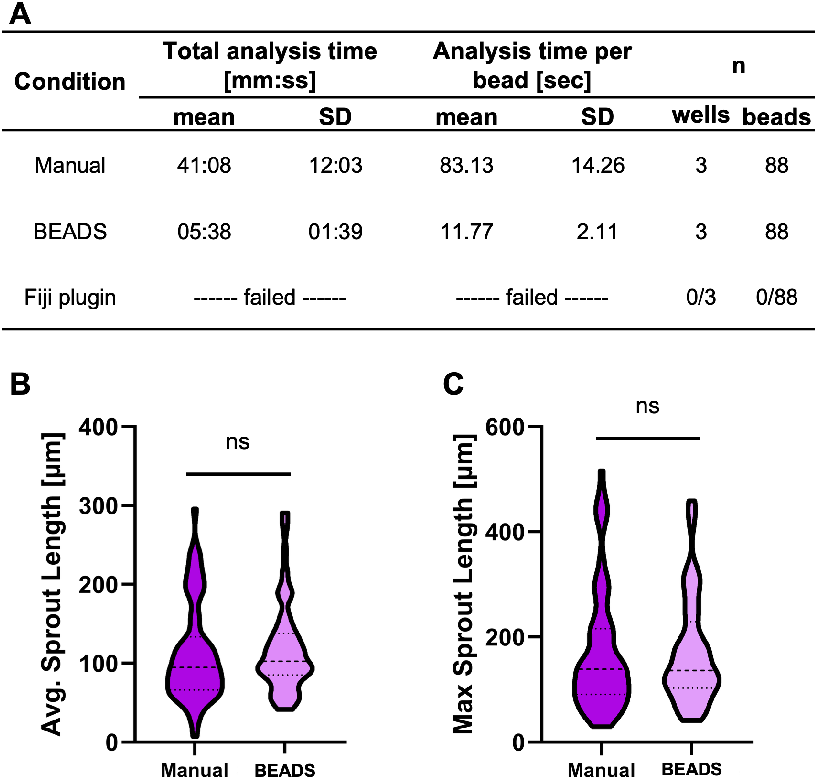
BEADS provides a sevenfold reduction in analysis time while preserving quantitative accuracy. **A)** Comparison of total analysis time per well and time per bead for manual tracing versus BEADS. The ImageJ Angiogenesis Analyzer (Fiji plugin) failed to identify beads or sprouts and therefore did not provide quantitative measurements. Average **(B)** and maximum **(C)** sprout length per bead measured by manual tracing and BEADS (n = 88 beads). Violin plots show mean ± IQR. Sprout-length measurements did not differ significantly between methods, confirming the accuracy of the BEADS workflow.

## DISCUSSION

The 3D fibrin bead assay remains a powerful tool for interrogating endothelial sprouting, migration, and microvascular morphogenesis; however, its broad adoption has been hindered by analysis workflows that are either slow, too labor-intensive, or insufficiently adaptable to heterogeneous imaging conditions. In this study, we developed BEADS, a semi-automated, user-interactive image-analysis pipeline designed to overcome these limitations and enable robust quantification of co-cultured endothelial populations, spatial relationships, and sprout-orientation behaviors.

A key design choice was to use semi-automation rather than full automation, allowing user corrections when heterogeneity, overlapping structures, or mosaic staining challenge segmentation algorithms. This hybrid approach prevents loss of datasets that would otherwise fail fully automated pipelines while reducing annotation time by roughly sevenfold. The balance between automation and expert oversight is particularly important in microvascular imaging, where typical fibrin bead images often exhibit signal heterogeneity, complex multicellular structures, and faint sprouts. This framework can thus standardize morphometric endpoints across microvascular laboratories, supporting reproducible quantification of angiogenic processes.

Using co-cultured male and female HPMECs, we showed that intrinsic sprouting metrics were not sex-dependent under hormone-free conditions, while female endothelial cells demonstrated significantly higher migratory behavior **(Fig 3)**. These findings are consistent with known sex-dependent endothelial phenotypes (16, 17) and confirm that the assay captures biologically relevant differences in motility. Importantly, the co-culture format ensures that both sexes experience identical biochemical and mechanical environments, enabling isolation of intrinsic, population-specific behaviors.

BEADS further enabled a novel nearest-neighbor spatial analysis, revealing that male bead-oriented sprouts preferentially oriented toward female neighbors **(Fig 4C)**, an effect that would be inaccessible using classical morphometric metrics. These results suggest sex-dependent local cues through paracrine or contact-dependent signaling influence microvascular alignment and highlight the importance of relational spatial metrics for understanding angiogenic patterning. While the mechanisms underlying these effects remain to be defined, the analytic capacity to detect them represents a substantial methodological advance.

This study presents a representative use case for the 3D fibrin angiogenesis assay by co-culturing M- and F-coated beads in the same well. However, BEADS is not limited to co-cultures of M and F cells. The platform is adaptable to a variety of image-based angiogenesis experiments, including different endothelial subtypes, disease models, pharmacologic perturbations, and other multicellular co-culture systems, provided that cell populations can be differentiated by labeling or marker expression (4, 33–36). In screening contexts, per-bead outputs (sprout presence, number, mean/max length, migration) can be aggregated to derive well-level dose-response metrics and hit calling, with the semi-automated workflow enabling higher-throughput experiments while preserving user oversight. It can also analyze single-cell populations. During analysis, users can select only one cell population by consistently selecting a single designation (“A” or “B”). Alternatively, these designations can represent additional bead characteristics such as donor, age, or cell subtype. In this study, membrane dyes were used to distinguish M- and F-cell populations prior to pooling them into a single well. While effective initially, membrane staining became less robust after three days in culture, making it difficult to differentiate the sex of some cells. Staining at the end of the assay with cell-type-specific markers for cell-type discrimination is a potential alternative; however, antibody staining within fibrin gels can be challenging. Future improvements could also integrate adaptive thresholding and machine-learning-based population classification to support multiplexed or multi-label microvascular assays.

## Conclusion

We demonstrate that images from 3D fibrin bead angiogenesis assays can be analyzed using a semi-automated, bead-centric approach implemented in BEADS. More specifically, BEADS enables quantification of sprout initiation, sprout length, migration, and nearest-neighbor directionality within co-cultured endothelial populations. Using mixed-sex HPMECs, we validated the platform’s accuracy, speed, and analytical breadth and revealed sex-dependent migratory behavior and directional sprouting toward female nearest neighbors. Representative datasets and annotated output files are available on the project’s GitHub repository to support on-boarding and validation by new users. These include raw image files of individual wells, bead/sprout annotations, and matched manual measurements for benchmarking. Together, these resources provide an accessible tool for standardized, reproducible microvascular image analysis, ensuring reproducibility and utility across laboratories. These capabilities position BEADS as an extensible platform for quantitative microvascular analysis and provide new opportunities to investigate endothelial interactions, co-culture dynamics, spatial pattern formation, and pharmacologic perturbation screens at the single-bead and population levels.

## ACKNOWLEDGEMENTS

The authors thank Katherine M. Nelson, Ph.D., for reviewing and commenting on the manuscript. This work was supported in part by the National Institutes of Health Grants F31HL152611 (BHP), T32GM133395 (BHP), R01DE029655(JPG), and R01HL144775 (JPG). Figures were created with a licensed version of Biorender.com

## DATA AND CODE AVAILABILITY

MATLAB Code is available at https://github.com/Gleghorn-Lab/BEADS_AngiogenesisWorkflow.

## AUTHOR CONTRIBUTIONS

Conceptualization (CRG, BHP, JPG), Methodology (CRG, JPG), Investigation (CRG, BHP), Data Analysis (CRG, JPG), Writing Original Draft (CRG), Manuscript Review Editing, & Approval (CRG, BHP, JPG), Supervision (JPG), Project Administration (JPG), Funding acquisition (BHP, JPG)

## Supplementary Information

### Protocol for running the Gleghorn Lab Bead-based Endothelial Angiogenesis Data Suite (BEADS) using MATLAB

#### Before you begin

1. Make sure MATLAB is installed on your computer
2. Download the “BEADS_AngiogenesisWorkflow” folder from GitHub
  a. https://github.com/Gleghorn-Lab/BEADS_AngiogenesisWorkflow
  b. Contents should include:
    - `AngiogenesisSproutingAssayBeadFindingCode.m`
    - `MigratingCellCounting.m`
    - `PostProcessing.m`
    - `README.md`
3. Download the Bio-Formats package for MATLAB https://docs.openmicroscopy.org/bio-formats/5.8.2/developers/matlab-dev.html
  a. Extract all files and copy them into the “BEADS_AngiogenesisWorkflow-main” folder

#### Bead sprouting analysis

1. Open the file labeled `AngiogenesisSproutingAssayBeadFindingCode.m` in MATLAB.
2. Before running the code, first ensure that the code is accurately assigned to the color channels of your image: 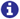 **NOTE:** If using `.CZI` file types (Zeiss proprietary format), they must have the brightfield image first, followed by the two fluorescent channels. If images are not acquired in this order, the order must be changed specifically in the MATLAB source code based on the channel order specified in Zen. The order of your colored channels is controlled by lines 35-38 and lines 130-133 of the code. Your channel order listed in lines 35-38 should always match that listed in 130-133.
  - Brightfield (BF) should always be assigned to variables “ch1” (line 36) and “ch1_2” (line131). The first numbers in the parentheses are dependent on the order of the laser channels in Zen. For example, if your channels appear in the order below,

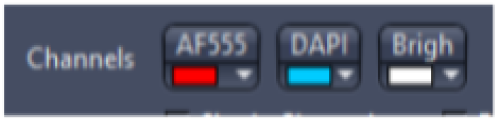 then your code should read:

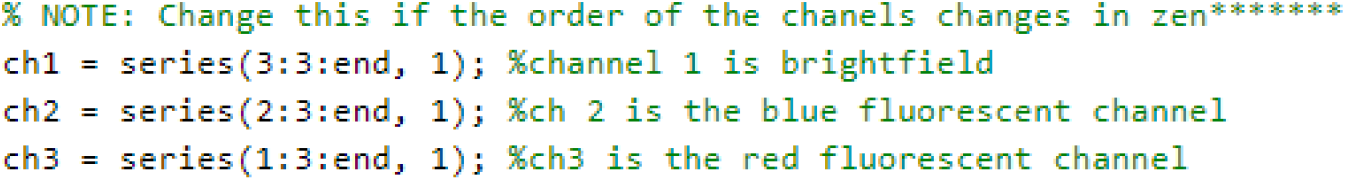 As the BF channel is listed 3rd in Zen, i.e., 3, the DAPI channel is listed 2nd in Zen, i.e., 2, and the AF555 channel is listed 1st in Zen, i.e., 1.
  - ONLY the very first number in the parentheses should change.
  - For example, if the brightfield is now the first channel in Zen and no longer the third channel listed in Zen, the numbers change from: ch1 = series(**3**:3:end, 1) → series(**1**:3:end, 1). ONLY the **BOLDED** number changes. 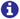 **NOTE:** If your image only has 2 channels, you’ll need to adjust the source code in more detail.
3. Once ensuring that the code is correct, press “Run”.
4. When prompted, select the desired .CZI file. The file should contain an image of the bead assay; this code has only been tested on 9×9-tiled 10x whole-well image z-stacks.
  a. Only one file may be selected at a time.
5. After selecting the desired image, a second window will appear, displaying all beads found by the code in GUI form.
  a. Any beads not detected can be selected by creating a circle around the bead. This can be done by clicking and dragging across the bead.
  b. To deselect a bead, right-click on the circle. 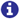 **NOTE:** The code will error in the next step if there is a bead < 200 µm from the edge of the image. To prevent this, you may deselect edge beads.
  c. Position and size of all circles can be adjusted by clicking and dragging on the ROI circle around the bead.
6. Once all beads have been selected, close the pop-up window by pressing the “x” in the top right corner. A new window will open, displaying the selected individual bead.
  a. The selected bead will have a yellow dot indicating its centroid. If there are other beads within the frame, you can ignore them, as the code stores data only for the bead with the yellow point at its center.
7. Click on the end of each sprout that is branching off the bead with the yellow center. You can right-click to deselect a sprout.
8. Once each sprout on the bead has been counted, press the “A” or “B” key on the keyboard to categorize the bead as belonging to population A or population B, respectively.
9. After designating the population, the next bead will be shown, and steps 7 and 8 should be repeated until all the beads have been shown
10. When the final bead has been assigned a population, the code will display an image of the entire well with red ‘*’ where all selected sprouts are. This is provided for confirmation only; if an error is detected at this step, the analysis must be restarted. Once done, the window can be closed.
11. Data should appear in the same folder as the MATLAB file that was run. There will be two text files that can be used in the PostProcessing code and an .xlsx file. The files will be named `BeadData.txt`, `SproutData.txt`, and `BeadAndSproutData.xlsx`. 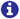 **NOTE: If the code is rerun without changing these file names, they will be overwritten, and data will be lost. Ensure that all output files are renamed**. Rename files to `NewFileName_BeadData.txt`, `NewFileName_SproutData.txt`, and `NewFileName_BeadAndSproutData.xlsx` 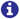 **NOTE: Do NOT** change the “SproutData” and “BeadData” portions of the .txt file names, as the post-processing code will use them for further analysis. Instead, add your new name as a prefix, separated by an underscore, as denoted above in step 11b.

#### Post processing

1. To obtain more detailed bead and sprout data in an Excel sheet, open the file labeled `PostProcessing.`m in MATLAB, and press run.
2. When prompted to select files, select the two .txt files that you relabeled in the previous step.
  a. You will first select the “BeadData” file, then the “SproutData” file.
  b. The code will process the data and analyze sprout length, bead spacing, the location of the nearest neighbor (NN), and the NN’s population type. All of this will be output in table form.
3. The new analyzed data file will appear in two formats `EndBeadSproutData.xlsx` and `EndBeadSproutData.txt` in the same folder as the MATLAB code.
  a. Rename the file, and transfer it to your own folder.
  b. If code is rerun without changing the file name, the file will be overwritten, and data will be lost. Ensure that all output files are renamed!

#### Migratory cell counting

1. Open the file labeled `MigratingCellCounting.m` in MATLAB.
2. Before running the code, first ensure that the code is accurately assigned to the color channels of your image. To change the order of your channels, follow instruction step 2 from the bead sprouting analysis instructions above.
3. Once ensuring that the code is correct, press “Run”.
4. When prompted, select the desired .CZI file. The file should contain an image of the bead assay. Only one file may be selected at a time. 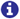 **NOTE:** this code has only been tested on 9×9 tiled 10x whole well image z stacks.
5. After selecting the desired image, a second window will appear. This window will be a small, cropped portion of the original image.
  a. The image will be cropped into an 81-tile grid of 9 rows x 9 columns.
  b. Cropped images will be shown on a per-row basis, always starting at column 1. So, the first window shown will be tile 1 – row 1 x column 1, and the 10th window shown will be tile 10 – row 2 x column 1.
6. Select all the migratory cells in the window by clicking on them. To deselect any cells, right-click on them.
7. To move on to the next window, press the “enter” key; there is no way to go back to a previous window. 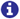 **NOTE: All 81 tiles will be cycled through TWO times**. The first cycle will allow you to select migratory cells from population A, and the second cycle will allow you to select cells from population B – “A” and “B” will be denoted at the top of the window, along with the tile number, to remind the user of where they are in the analysis process.
8. Data should appear in the same folder as the MATLAB file that was run and will be titled MigratoryCellCount.xlsx. 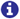 **NOTE: If the code is rerun without changing these file names, they will be overwritten, and data will be lost. Ensure that all output files are renamed**.

## Notes

https://github.com/Gleghorn-Lab/BEADS_AngiogenesisWorkflow

### Competing Interest Statement

The authors have declared no competing interest.

## References

1. Peter Carmeliet. Angiogenesis in health and disease. Nature medicine, 9(6): 653–660, 2003. ISSN 1078-8956.

2. Joshua T. Morgan, Wade G. Stewart, Robert A. McKee, and Jason P. Gleghorn. The mechanosensitive ion channel TRPV4 is a regulator of lung development and pulmonary vasculature stabilization. Cellular and Molecular Bioengineering, 11(5):309–320, . ISSN 1865-5033. doi: 10.1007/s12195-018-0538-7.

3. Brea Chernokal, Cailin R. Gonyea, and Jason P. Gleghorn. Lung development in a dish: Models to interrogate the cellular niche and the role of mechanical forces in development. In Chelsea M. Magin, editor, Engineering Translational Models of Lung Homeostasis and Disease, Advances in Experimental Medicine and Biology, pages 29–48. Springer International Publishing, . ISBN 978-3-031-26625-6. doi: 10.1007/978-3-031-26625-6_3.

4. Celeste M. Nelson and Jason P. Gleghorn. Sculpting organs: Mechanical regulation of tissue development. Annual Review of Biomedical Engineering, 14(1):129–154. ISSN 1523-9829, 1545-4274. doi: 10.1146/annurev-bioeng-071811-150043.

5. Jasmine Shirazi, Joshua T. Morgan, Erica M. Comber, and Jason P. Gleghorn. Generation and morphological quantification of large scale, three-dimensional, self-assembled vascular networks. MethodsX, 6:1907–1918. ISSN 2215-0161. doi: 10.1016/j.mex.2019.08.006.

6. Katherine M Nelson, N’Dea Irvin-Choy, Matthew K Hoffman, Jason P Gleghorn, and Emily S Day. Diseases and conditions that impact maternal and fetal health and the potential for nanomedicine therapies. Advanced drug delivery reviews, 170:425–438, . ISSN 1872-8294. doi: 10.1016/j.addr.2020.09.013. Edition: 2020/09/28 Place: Netherlands.

7. Jason P. Gleghorn and Megan L. Killian. Chapter 3 - mechanobiology throughout development. In Stefaan W. Verbruggen, editor, Mechanobiology in Health and Disease, pages 77–98. Academic Press. ISBN 978-0-12-812952-4. doi: 10.1016/B978-0-12-812952-4.00003-9.

8. Gabriele Bergers and Laura E Benjamin. Tumorigenesis and the angiogenic switch. Nature reviews cancer, 3(6):401–410, 2003. ISSN 1474-175X.

9. Hanna M Eilken and Ralf H Adams. Dynamics of endothelial cell behavior in sprouting angiogenesis. Current opinion in cell biology, 22(5):617–625, 2010. ISSN 0955-0674.

10. Samuel I. Hofbauer, Luisa A. Fink, Rachel E. Young, Tara Vijayakumar, Katherine M. Nelson, Nia Bellopede, Mohamad-Gabriel Alameh, Drew Weissman, Jason P. Gleghorn, and Rachel S. Riley. Cytokine mRNA delivery and local immunomodulation in the placenta using lipid nanoparticles. Advanced Therapeutics, 8(8):e00148. ISSN 2366-3987. doi: 10.1002/adtp.202500148.

11. Majid Sharifi, William C. Cho, Asal Ansariesfahani, Rahil Tarharoudi, Hedyeh Malekisarvar, Soyar Sari, Samir Haj Bloukh, Zehra Edis, Mohamadreza Amin, Jason P. Gleghorn, Timo L. M. ten Hagen, and Mojtaba Falahati. An updated review on EPR-based solid tumor targeting nanocarriers for cancer treatment. Cancers, 14(12):2868. ISSN 2072-6694. doi: 10.3390/cancers14122868. Number: 12 Publisher: Multidisciplinary Digital Publishing Institute.

12. N’Dea S Irvin-Choy, Katherine M Nelson, Jason P Gleghorn, and Emily S Day. Design of nanomaterials for applications in maternal/fetal medicine. Journal of materials chemistry. B, 8(31):6548–6561. ISSN 2050-7518. doi: 10.1039/d0tb00612b. Edition: 2020/05/26 Place: England.

13. Julia Felsenstein, Katherine M. Nelson, Jasmine Shirazi, and Jason P. Gleghorn. Therapeutic nanoparticle safety in pregnancy: Bridging knowledge gaps with environmental insights and a translational roadmap. Journal of Controlled Release, page 114026. ISSN 0168-3659. doi: 10.1016/j.jconrel.2025.114026.

14. Volker Nehls and Detlev Drenckhahn. A Novel, Microcarrier-Based in Vitro Assay for Rapid and Reliable Quantification of Three-Dimensional Cell Migration and Angiogenesis. Microvascular Research, 50(3):311–322, 1995. ISSN 0026-2862. doi: 10.1006/mvre.1995.1061.

15. Bruno Vailhé, Daniel Vittet, and Jean-Jacques Feige. In Vitro Models of Vasculogenesis and Angiogenesis. Laboratory Investigation, 81(4):439–452, 2001. ISSN 1530-0307. doi: 10.1038/labinvest.3780252.

16. Yuhao Zhang, Xiaoyu Dong, Jasmine Shirazi, Jason P Gleghorn, and Krithika Lingappan. Pulmonary endothelial cells exhibit sexual dimorphism in their response to hyperoxia. American Journal of Physiology-Heart and Circulatory Physiology, 315(5):H1287–H1292, 2018. ISSN 0363-6135.

17. Brielle Hayward-Piatkovskyi, Cailin R Gonyea, Sienna C Pyle, Krithika Lingappan, and Jason P Gleghorn. Sex-related external factors influence pulmonary vascular angiogenesis in a sex-dependent manner. American Journal of Physiology-Heart and Circulatory Physiology, 324(1):H26–H32, nov 2022. ISSN 0363-6135. doi: 10.1152/ajpheart.00552.2022.

18. Michael J. Donzanti, Bryan J. Ferrick, Omkar Mhatre, Brea Chernokal, Diana C. Renteria, and Jason P. Gleghorn. Stochastic to deterministic: A straightforward approach to create serially perfusable multiscale capillary beds. ACS Biomaterials Science & Engineering, 11(1):239–248. doi: 10.1021/acsbiomaterials.4c01247. Publisher: American Chemical Society.

19. Vani Narayanan, Laurel E. Schappell, Carl R. Mayer, Ashley A. Duke, Travis J. Armiger, Paul T. Arsenovic, Abhinav Mohan, Kris N. Dahl, Jason P. Gleghorn, and Daniel E. Conway. Osmotic gradients in epithelial acini increase mechanical tension across e-cadherin, drive morphogenesis, and maintain homeostasis. Current biology: CB, 30(4):624–633.e4. ISSN 1879-0445. doi: 10.1016/j.cub.2019.12.025.

20. Laurel E. Schappell, Daniel J. Minahan, and Jason P. Gleghorn. A microfluidic system to measure neonatal lung compliance over late stage development as a functional measure of lung tissue mechanics. Journal of Biomechanical Engineering, 142(100803). ISSN 0148-0731. doi: 10.1115/1.4047133.

21. Shelby R. Mohr-Allen, Jason P. Gleghorn, and Victor D. Varner. Fluid secretion and luminal pressure control lateral branching morphogenesis in the embryonic avian lung. Developmental Biology, 520:251–263. ISSN 1095-564X. doi: 10.1016/j.ydbio.2025.01.016.

22. Kara E. Garcia, Wade G. Stewart, M. Gabriela Espinosa, Jason P. Gleghorn, and Larry A. Taber. Molecular and mechanical signals determine morphogenesis of the cerebral hemispheres in the chicken embryo. Development, 146 (20):dev174318. ISSN 1477-9129, 0950-1991. doi: 10.1242/dev.174318.

23. Catherine S. Millar-Haskell, John L. Sperduto, John H. Slater, Colin Thorpe, and Jason P. Gleghorn. Secretion of the disulphide bond generating catalyst QSOX1 from pancreatic tumour cells into the extracellular matrix: Association with extracellular vesicles and matrix proteins. Journal of Extracellular Biology, 1(7):e48,. ISSN 2768-2811. doi: 10.1002/jex2.48. _eprint: https://onlinelibrary.wiley.com/doi/pdf/10.1002/jex2.48.

24. Yuhao Zhang, Cristian Coarfa, Xiaoyu Dong, Weiwu Jiang, Brielle Hayward-Piatkovskyi, Jason P Gleghorn, and Krithika Lingappan. MicroRNA-30a as a candidate underlying sex-specific differences in neonatal hyperoxic lung injury: implications for BPD. American Journal of Physiology-Lung Cellular and Molecular Physiology, 316(1):L144–L156, nov 2018. ISSN 1040-0605. doi: 10.1152/ajplung.00372.2018.

25. Philipp Berens. CircStat: A MATLAB Toolbox for Circular Statistics. Journal of Statistical Software, 31(10 SE - Articles):1–21, sep 2009. doi: 10.18637/jss.v031.i10.

26. Landler Lukas, Graeme D Ruxton, and Malkemper E Pascal. Grouped circular data in biology: advice for effectively implementing statistical procedures. Behavioral Ecology and Sociobiology, 74(8), aug 2020. ISSN 0340-5443. doi: 10.1007/s00265-020-02881-6.

27. Joshua T. Morgan, Jasmine Shirazi, Erica M. Comber, Christian Eschenburg, and Jason P. Gleghorn. Fabrication of centimeter-scale and geometrically arbitrary vascular networks using in vitro self-assembly. Biomaterials, 189: 37–47,. ISSN 01429612. doi: 10.1016/j.biomaterials.2018.10.021.

28. Krithika Lingappan, Oluyinka O. Olutoye, Abiud Cantu, Manuel Eliezer Cantu Gutierrez, Nahir Cortes-Santiago, J. D. Hammond, Jamie Gilley, Joselyn Rojas Quintero, Hui Li, Francesca Polverino, Jason P. Gleghorn, and Sundeep G. Keswani. Molecular insights using spatial transcriptomics of the distal lung in congenital diaphragmatic hernia. American Journal of Physiology-Lung Cellular and Molecular Physiology, 325(4):L477–L486. ISSN 1040-0605. doi: 10.1152/ajplung.00154.2023. Publisher: American Physiological Society.

29. Cailin R. Gonyea, Yuanjun Shen, Katherine M. Nelson, Rylie N. Bird, Rachel M. Gilbert, Oluyinka O. Olutoye, Sundeep G. Keswani, and Jason P. Gleghorn. The nitrofen/bisdiamine murine model of congenital diaphragmatic hernia has a pulmonary hypertension vascular phenotype consistent with human CDH. American Journal of Physiology. Lung Cellular and Molecular Physiology. ISSN 1522-1504. doi: 10.1152/ajplung.00233.2024.

30. Virginia H Huxley, Scott S Kemp, Christine Schramm, Steve Sieveking, Susan Bingaman, Yang Yu, Isabella Zaniletti, Kevin Stockard, and Jianjie Wang. Sex differences influencing micro-and macrovascular endothelial phenotype in vitro. The Journal of physiology, 596(17):3929–3949, 2018.

31. Anna E Stanhewicz, Megan M Wenner, and Nina S Stachenfeld. Sex differences in endothelial function important to vascular health and overall cardiovascular disease risk across the lifespan. American Journal of Physiology-Heart and Circulatory Physiology, 315(6):H1569–H1588, 2018.

32. Felipe Troncoso, Kurt Herlitz, Jesenia Acurio, Claudio Aguayo, Katherine Guevara, Fidel Ovidio Castro, Alejandro S Godoy, Sebastian San Martin, and Carlos Escudero. Advantages in wound healing process in female mice require upregulation a2a-mediated angiogenesis under the stimulation of 17β-estradiol. International Journal of Molecular Sciences, 21(19):7145, 2020.

33. Katherine M. Nelson, Daniel J. Minahan, Vonetta L. Edwards, Ian J. Glomski, David J. Delgado Diaz, Keena Thomas, Forrest C. Walker, Patrik M. Bavoil, Isabelle Derré, Alison K. Criss, Jacques Ravel, and Jason P. Gleghorn. A microphysiologic model of the cervical epithelium recapitulates microbial, immunologic, and pathogenic properties of sexually transmitted infections. bioRxiv, . doi: 10.1101/2025.07.21.665989.

34. Catherine S. Millar-Haskell, Allyson M. Dang, and Jason P. Gleghorn. Coupling synthetic biology and programmable materials to construct complex tissue ecosystems. MRS communications, 9(2):421–432, . ISSN 2159-6859. doi: 10.1557/mrc.2019.69.

35. Brea Chernokal, Bryan J. Ferrick, and Jason P. Gleghorn. Zonal patterning of extracellular matrix and stromal cell populations along a perfusable cellular microchannel. Lab on a Chip, 24(23):5238–5250, . ISSN 1473-0189. doi: 10.1039/D4LC00579A. Publisher: The Royal Society of Chemistry.

36. Vonetta L Edwards, Elias McComb, Jason P Gleghorn, Larry Forney, Patrik M Bavoil, and Jacques Ravel. Three-dimensional models of the cervicovaginal epithelia to study host–microbiome interactions and sexually transmitted infections. Pathogens and Disease, 80(1):ftac026. ISSN 2049-632X. doi: 10.1093/femspd/ftac026.

